# Sleep immediately after noise overexposure worsens tinnitus in mice

**DOI:** 10.64898/2026.06.01.729258

**Authors:** Linus Milinski, Fernando R. Nodal, Andrew J. King, Vladyslav V. Vyazovskiy, Victoria M. Bajo

**Author notes:** Corresponding authors: LM and VMB.

## Abstract

Subjective tinnitus is the most common auditory phantom perception in the absence of an acoustic stimulus yet there is no cure nor widely effective treatment to date. Previously, we suggested that sleep may contribute to tinnitus development after a triggering event, such as noise overexposure (NOE).

Here, we investigated the effect of natural sleep on tinnitus development following NOE in mice. One group of mice was kept awake for 6 hours immediately after NOE, whereas a control group could sleep *ad libitum*. Tinnitus and hearing loss were assessed before and after NOE by determining gap prepulse inhibition of the acoustic startle reflex (GPIAS) and by recording auditory brainstem responses (ABRs).

Only mice that were allowed to sleep in the first six hours after NOE developed a significant change in the GPIAS ratio one week after NOE, indicative of tinnitus. Habituation to the startle stimulus was largely similar between both groups, except that at eight weeks after NOE, when GPIAS ratio had returned to baseline levels in both groups, the sleep deprived group showed less habituation to the startle stimulus.

Notably, animals in neither group showed significantly elevated ABR thresholds after NOE, yet the control group showed signs of elevated evoked activity in the auditory brainstem one week after NOE.

These findings demonstrate that sleep early after noise overexposure may amplify subsequent tinnitus development. Our results suggest that sleep represents a temporal window that may be harnessed for potential modulation or mitigation of noise-induced hearing loss and tinnitus-related impairments.

## Introduction

Subjective tinnitus, a persistent phantom auditory perception that affects approximately 15% of the global population and currently lacks an effective cure, is often triggered by noise trauma. Yet it is now widely acknowledged that a potential tinnitus trigger event alone is not sufficient for the development of a persistent phantom percept ^1^. In fact, tinnitus development is considered to consist of two stages: first, a triggering event and its immediate effects on the integrity of the auditory system and, second, the subsequent consolidation of structural and functional changes in the brain, which are ultimately responsible for the persistent tinnitus ^2^. Within the central auditory brain these changes likely entail enhancement of sensory gain to compensate for loss of afferent inputs and maintain firing homeostasis ^3,4^, which may lead to elevated firing and bursting activity in various regions across the auditory pathway [dorsal cochlear nucleus ^5–8^; inferior colliculus ^9,10^; medial geniculate body ^11^; auditory cortex ^12–14^]. Such changes are especially prominent in the dorsal cochlear nucleus (DCN), where increased neuronal synchrony can be observed in animal models of tinnitus even without auditory threshold changes ^15^. This has been related to activity in DCN fusiform cells, which are, due to their multisensory inputs, particularly sensitive to spike-time-dependent plasticity.

Importantly, neural plasticity during tinnitus is not restricted to the auditory system but involves a change in activity across cortical hemispheres, including prefrontal cortex, orbitofrontal cortex and the parieto-occipital region ^16^. It has been suggested that tinnitus only emerges once brain-wide networks, consisting of frontal and parietal areas, are affected ^17,18^. Of particular importance are changes in limbic regions, such as emergence of neuronal hyperactivity in nucleus accumbens (NAc) ^19^ or a reduction of amygdala activation by unpleasant sounds, possibly due to an ‘internal modification of the emotional response’ ^20^. Such limbic involvement in tinnitus has been related to the emotional impact of the phantom sound ^17,21,22^. Limbic dysregulation may also be a major factor in driving distress associated with tinnitus ^23^. In addition, brain-wide tinnitus representation that includes limbic areas may indicate tinnitus consolidation ^17,21,22^.

It is likely therefore that local and global brain plasticity are associated with tinnitus emergence and may, in addition, be instrumental in mediating generation of the persistent phantom percept. Indeed, electrical stimulation of the left vagus nerve paired with auditory stimulation can reverse perceptual impairments associated with noise-induced tinnitus in a rat model ^12^. In humans, vagus nerve stimulation can lead to small but clinically significant improvements in tinnitus loudness and distress ^24^, suggesting that brain-triggered plasticity could be the basis for a treatment. Yet, so far, the possibilities for exploiting brain plasticity to understand and treat tinnitus are restricted by a lack of understanding of its scope and timescale after the tinnitus trigger. One intriguing way to address this is to focus on naturally occurring time windows when the brain is susceptible to plastic changes and harness this potential for experimental manipulation and possible new routes of treating tinnitus.

Each day, the brain cycles through functional states based on different global spatiotemporal activity patterns, especially across wakefulness and sleep ^25^. Sleep-wake dynamics are shaped by the endogenous circadian clock and the light-dark cycle, as well as by homeostatic (sleep-wake history) factors ^26,27^. Wakefulness involves both sensory-evoked and spontaneous activity, whereas spatiotemporal brain dynamics during sleep are primarily driven by spontaneous activity. While neural activity during wakefulness promotes neural plasticity, the sleep state is thought to be of particular importance in consolidating plastic changes ^28–33^.

Consolidation of experience-dependent plasticity in particular is thought to rely on sleep-dependent mechanisms, such as synaptic strengthening and weakening ^34–36^ possibly based on the replay of memory traces ^31^. It has been shown that cortical regions involved in executing tasks during wakefulness exhibit more pronounced slow-wave activity during subsequent sleep ^37,38^. In turn, if slow waves are suppressed or disrupted, this can interfere with the benefits sleep might otherwise provide for memory formation. Slow-waves during NREM sleep have been proposed to be instrumental in sleep-dependent downscaling of synaptic strength, a process thought to be essential for sustaining brain plasticity ^39^. Furthermore, during NREM sleep, depolarisation of a local cortical cell population can increase the chance of concerted neural firing and, in turn, promote plasticity ^40^.

The possibility remains, therefore, that plasticity in the auditory system, subsequent to noise trauma or a reduction of peripheral input, is also a state-dependent process and that neural plasticity during NREM sleep might be a contributor to tinnitus formation ^41^. Even though plasticity observed during sleep shows similarities to plasticity thought to underly tinnitus, the role of sleep in tinnitus development remains unclear. Here, the effects of sleep and sleep-deprivation on tinnitus development were evaluated in a mouse model of noise-induced tinnitus.

## Methods

### Animals and experimental design

Adult male C57/BL6ntac-cdh23h-9 mice (n=14) were housed on a 12/12 light-dark cycle. Food and water were available *ad libitum*. The C57/BL6ntac-cdh23h-9 strain was chosen for this study as these animals, due to a corrected Cdh23(ahl) allele, do not show early-onset age-related hearing loss that is present in commonly used inbred mouse strains ^42^.

All animals were implanted with frontal and occipital EEG screw electrodes and neck muscle electrodes at the beginning of the experiment. After at least a week of recovery, baseline gap prepulse inhibition of the acoustic startle reflex (GPIAS) and auditory brainstem responses (ABRs) were assessed in all animals (Fig. 1).

**Figure 1.**
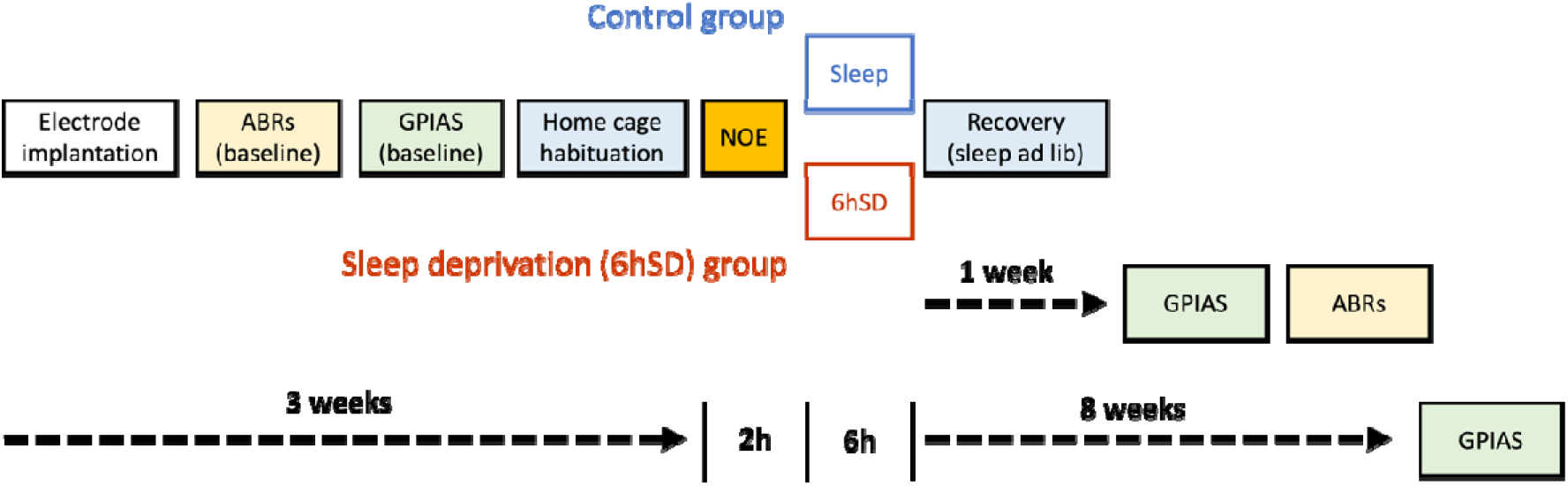
Outline of the experiment. Animals were implanted with frontal and occipital EEG screw electrodes and neck muscle electrodes at the beginning of the experiment. After recovery, auditory brainstem responses (ABRs) were assessed in all animals. Subsequently, baseline gap prepulse inhibition of inhibition of the acoustic startle (GPIAS) was tested. Mice were then transferred into new home cages suitable for video and EEG monitoring. After a minimum of four days of habituation to the new environment and tethering, noise overexposure (NOE, 8kHz centred NBN presented at 98dB SPL) was administered under anaesthesia for a duration of 2 hours at the end of the dark phase. Immediately afterwards, at light onset, animals were transferred to a heated recovery chamber for up to 5 minutes until recovery from overt anaesthesia. Mice were then transferred back to their home cages. Animals in the 6hSD group (n=8) were kept awake for a period of six hours while EEG signals were monitored. Animals in the control group (n=6) were left undisturbed after experiencing NOE. About 1 week (ca 6 days) after NOE, animals where again assessed in the GPIAS paradigm and, subsequently, ABR responses were obtained. About 8 weeks after noise overexposure, GPIAS was assessed for a third time.

Mice were then transferred to recording cages (suitable for obtaining video and EEG recordings in the freely-moving animals), which also served as home cages. After at least four days of habituation to the new cage and being tethered to the EEG recording setup, they were exposed to noise under anaesthesia for 2 hours at the end of the dark phase (ZT22–24) (Fig. 1). Immediately afterwards, at light onset, mice were transferred to a heated (25-28 °C) recovery chamber for up to 5 minutes until the animals recovered from overt anaesthesia (i.e. displayed spontaneous locomotion). Mice were then transferred back to their home/recording cages.

Animals in the sleep deprivation (6HSD) group (n=8) were kept awake for a period of six hours. Animals in the control group (n=6) were left undisturbed after having been transferred back to their home/recording cages and could sleep *ad libitum* after experiencing NOE. The time of NOE (at the end of the dark period, the animals’ active phase) was chosen to ensure that sleep deprivation would occur during the animals’ natural sleep time (within the first half of the light phase) and thus ensure that the difference in sleep amount between control and six hours sleep deprived (6hSD) animals would be maximal.

Approximately one week after NOE (Post1w), animals were again assessed in the GPIAS paradigm and ABRs were recorded. GPIAS was assessed for a third time about eight weeks after NOE (Post8w) (see Fig. 1 for a timeline of the experiment) based on previous research in mice describing chronic tinnitus 5-7 weeks after noise overexposure [see Fig. 4 in Turner et al 2012 ^43^). Note that given the extended timeline of the experimental protocol, including repeated anaesthesia for ABR and surgical procedures, ABRs were acquired at only one time point after NOE in adherence to the 3Rs principles.

### Implantation of electrodes

The surgical procedure largely followed previously described protocols [e.g 42,43]. Electrode implantation was carried out in sterile conditions under isoflurane anaesthesia (3–5% induction, 1–2% maintenance). During surgery, animals were head-fixed using a stereotaxic frame (David Kopf Instruments, CA, USA). Liquid gel (Viscotears, Alcon Laboratories Ltd., UK) was applied to protect the eyes. Metacam (5mg/kg, s.c.) and Vetergesic (0.1 mg/kg s.c.) were administered preoperatively to reduce inflammation and discomfort.

Stainless screw electrodes were placed unilaterally in the left frontal (AP +2 mm, ML - 2 mm) and occipital (AP −3.5 to −4 mm, ML - 2.5 mm) cortical regions. A reference screw electrode was placed above the cerebellum. EEG screws were soldered (prior to implantation) to custom-made headmounts (Pinnacle Technology Inc. Lawrence, USA), and all screws and wires were secured to the skull using dental acrylic. Two single-stranded stainless-steel wires were inserted on either side of the nuchal muscle to record the electromyogram (EMG). Screw-electrode placement optimised detection of frontal slow-wave and occipital theta activity for vigilance-state assessment. In this experiment, the screw electrodes were used only for vigilance state monitoring, not for assessment of auditory function. Auditory function was assessed via ABR recordings using separate, acutely placed subcutaneous electrodes that specifically recorded auditory brainstem-evoked activity (see Methods section: Auditory brainstem response (ABR) measurements).

At the end of the surgery, animals were administered saline (0.1 ml/20 g body weight, s.c.). The animal’s body temperature was monitored and maintained throughout the procedure using a rectal probe and a homeothermic monitoring system (Harvard Apparatus), and a heated chamber for recovery afterwards until the animal readily walked around. Metacam (1–2 mg/kg) was orally administered for 3 days after surgery. A minimum 2-week recovery period was provided prior to further procedures.

### Noise overexposure (NOE)

To trigger tinnitus, all mice were temporarily exposed to loud noise (noise overexposure, NOE), as is commonly used in animal models of tinnitus ^6,9,12,43^, including in mice ^43^.

Noise (1/2 octave narrowband pink noise centred at 8 kHz, 98 dB SPL at ear level) was presented for 120 minutes via a free-field loudspeaker positioned approximately 8 cm in front of the animal and facing it. The left ear was protected by fitting it with an earplug and silicone impression material (Otoform, Dreve). The procedure was carried out under isoflurane anaesthesia (3–5% induction, 1–2% maintenance). Liquid gel (Viscotears, Alcon Laboratories Ltd., UK) and patches of aluminium foil were applied to protect the eyes and shield from light exposure. The animal’s temperature was monitored and maintained throughout the procedure using a rectal probe and a homeothermic monitoring system (Harvard Apparatus), and a heated chamber for recovery afterwards until the animal readily walked around. NOE took place during the last two hours of the dark phase (ZT22-24) for all animals.

### Tinnitus assessment

Tinnitus was assessed in a variant of behavioural gap-in-noise detection (GPIAS), which is commonly used to assess tinnitus in the mouse model ^43,46^.

The animals were placed in a small enclosure (Fig.2A) positioned on a motion-sensitive platform (San Diego Instruments, SR-LAB Startle Response System) and presented with a white noise acoustic background at 67 dB SPL, similar to the paradigm used by Turner et al. 2012 ^43^. Each session began with an acclimatisation period that consisted of a 3 minutes with only background noise, followed by two startle-eliciting white noise bursts (20 ms, 105 dB SPL), to obtain a more stable baseline response ^43^. Data from the acclimatisation period were not used in subsequent analysis.

**Figure 2.**
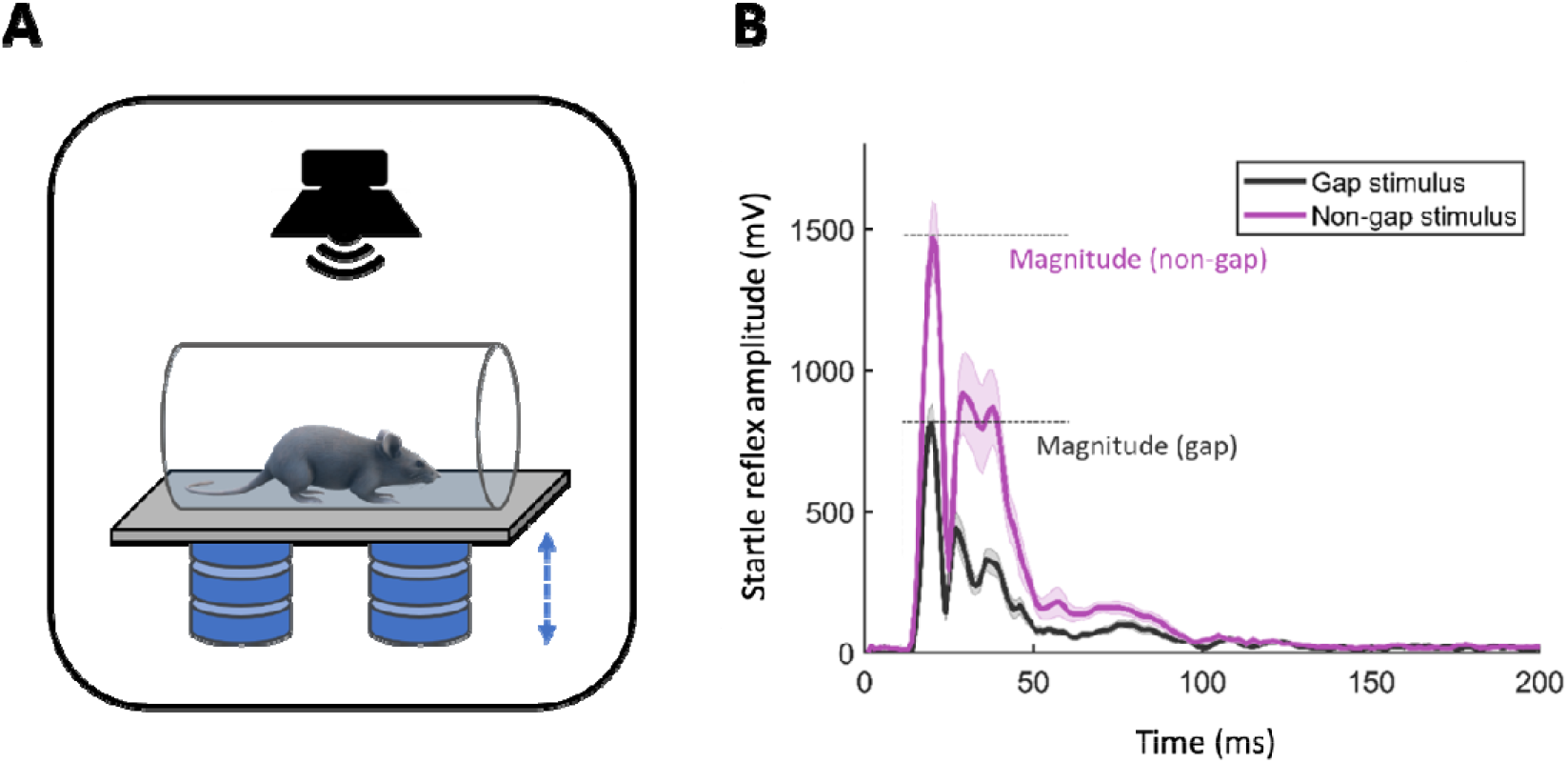
Gap pre-pulse inhibition of the acoustic startle (GPIAS). **(A)** Schematic depiction of the setup used for the GPIAS paradigm. Animals were placed in a tube-shaped chamber, which was positioned on a motion sensitive platform. Sound stimuli were presented via a loudspeaker located above the chamber. The chamber and loudspeaker were housed in an enclosure (not shown). **(B)** Example of an average startle signal in gap trials (black) and non-gap trials (purple). Typically, the startle reflex amplitude is higher in non-gap trials than in trials where the startle sound is preceded by a predictive gap in the ongoing background noise. The startle response was assessed in two ways: first, by measuring the magnitude of only the most pronounced peak in the signal (defined as maximum - minimum value of the signal within the response window, highlighted by horizontal dashed lines in the figure) and, second, by calculating the average of the signal within the response window. These measures were assessed for the onset of the startle response (initial 50ms of the signal), or over the entire recorded response (response window of 1000ms). If not stated otherwise, reported GPIAS values are based on combined data based on both measures (peak signal and average of the signal) and response windows (0-50s or 0-1000ms).

After acclimatisation, 40 startle-eliciting noise bursts were presented at random intervals (3-13 s) over the background noise. Half of the startle stimuli were preceded by a silent gap (50ms gap duration) in the background noise 100 ms before the startle noise. This commonly leads to a reduced startle response relative to that evoked by startle stimuli that are not preceded by a silent gap. This decrease in the amplitude of the startle response is referred to as GPIAS (gap-induced prepulse inhibition of the acoustic startle) and is thought to provide a measure of gap detection ability ^43,47^ (Fig.2B). Since animals are prone to habituation in this paradigm (i.e., a reduced startle response after repeated exposure to the startle stimulus), each assessment consisted of only 40 trials (20 trials with a silent gap preceding the startle stimulus and 20 trials without; trials were randomly interspersed). In addition, consecutive GPIAS assessments were separated by at least 14 days.

The startle amplitude was calculated for each trial offline. For every trial, two parameters were used to quantify the startle response. First, the peak amplitude, defined as maximum - minimum value of the signal within the response window. The second parameter was the average of the signal within the response window. Both measures were assessed in a narrow and a wider response window: one to capture the initial startle response (in the first 50ms after presentation of the startle stimulus), and the other over one second (within the first 1000ms after presentation of the startle stimulus) to capture temporal dynamics in the startle response not reflected in the initial load cell deflection. Unless specified otherwise, the different metrics were assessed and combined for the analysis to capture multiple aspects of the startle response and hence reduce the impact of inter-trial variability in any one metric.

Gap detection ability was defined as the ratio between startle responses in gap trials and non-gap trials.

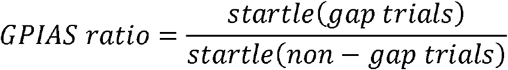

The GPIAS ratio was assessed in sequential blocks of five gap and five non-gap trials, providing measures for the first, second, third, and fourth quarters of a session. This block-based analysis has been previously suggested to capture effects of within-session habituation ^48^. Extreme startle response values lying outside the mean ± 5 standard deviations (based on a 5000-iteration bootstrap) were excluded from the analysis.

### Auditory brainstem response (ABR) measurements

To assess noise-induced hearing impairment ^43,49^, auditory brainstem responses (ABRs) were recorded before and after NOE under anaesthesia induced with i.p. medetomidine hydrochloride (0.022 mg/kg body weight; Domitor, Orion Pharma) and ketamine hydrochloride (5 mg/kg; Narketan10, Vetoquinol). Depth of anaesthesia and respiratory rate were monitored throughout the experiment via a camera focused on the animal. Animal temperature was monitored and maintained using a rectal probe and a homeothermic monitoring system (Harvard Apparatus). To expedite recovery from anaesthesia after the end of the procedure, animals were administered Antisedan (atipamezole hydrochloride, 0.06mg/kg, i.p.).

ABR signals were recorded from sterile subcutaneous monopolar needle electrodes (0.35 x 12 mm, MN3512P150, Spes Medica) placed close to the right and left bullae, and referenced to an electrode placed at the vertex of the skull. A ground electrode was positioned subcutaneously on the back of the animal. Signals were routed to a preamplifier (Medusa RA16PA Tucker Davies Technologies (TDT)) and recorded by an RZ2 Bioamp Processor controlled by BioSigRP software (TDT).

Sound stimuli were presented to one ear each time, while the other ear was fitted with an ear plug. Stimuli were presented from a free-field loudspeaker positioned 8 cm in front of the animal. Auditory stimuli were generated using an RP2.1 Enhanced Real-time processor (TDT), with a sampling frequency of 100 kHz, connected to a TDT PA5 programmable attenuator. The loudspeaker was calibrated using SigCalRP TDT calibration software to generate compensation filters, ensuring stable output levels for a frequency range from 250 to 30,000 Hz. Click stimuli were presented at a rate of 17/sec for 700 repetitions per level (40, 50, 60, 70, 80, 90 dB SPL). Pure tone stimuli (1, 4, 8 and 16 kHz) of 5 ms duration were presented at a rate of 21/sec for 700 repetitions per level and frequency combination.

### ABR threshold detection

Data analysis was performed offline based on the average ABR signals (averaged over 700 individual ABR traces) for each stimulus type. Data shown are based on ABRs of the right ear, if not specified otherwise. ABR thresholds were determined using the protocol described in Milinski et al., 2024 ^50^. Thresholds were determined manually by an experienced experimenter through visual assessment of ABR traces. This was conducted under blind conditions (enabled through a randomisation process used to access the data) with respect to animal, stimulus and stage of the experimental timeline (baseline, one week after NOE). Thresholds were defined as the lowest stimulus level where an ABR wave was present if corresponding waves were also present at higher sound levels. If no ABR wave was present for any sound level, the threshold was defined to be at 90 dB SPL (the highest sound level used).

### Automated ABR signal detection

Data analysis was performed offline based on average signals (averages over 700 individual ABR traces) for each stimulus type and level using a custom-written MATLAB script: local maxima and minima in the signal were detected using the *findpeaks* function (signal processing toolbox, MATLAB) and ABR waves I to V were defined depending on predetermined response windows for each sound level (40-90 dB SPL).

Waves were always detected first for the 90 dB SPL signal for any given stimulus (clicks or NBN stimuli) and following previous descriptions ^51–53^. The findpeaks function was applied to the entire signal. During the baseline condition (BL), the peak of the consistently most prominent wave, wave II was defined as the maximum peak between (and including) 1.6ms and 2.6ms. Peaks of waves I, III, IV and V were selected relative to the latency of wave II with the following criteria: −1.08ms ≤ peak I ≤ −0.4ms; 0.04 ≤ peak III ≤ 0.92 ms; 0.96ms ≤ peak IV ≤ 1.52ms; 1.56ms ≤ peak V ≤ 2.2ms. As wave latencies increase with decreasing sound level, the detection windows for sound levels below 90 dB SPL were shifted (delayed) by 0.16ms/10 dB decrement. Artefactual bias was minimised by excluding detected peaks: (1) if there was no detected peak for the next higher sound level (e.g., a wave II at 40 dB SPL was deemed artefactual if wave II was not detectable at 50 dB SPL) or (2) if a detected peak had a shorter latency than the corresponding peak at the next higher level with a tolerance of ±0.12 ms (3 data points, dps).

After detection of waves I to V, wave magnitudes were defined as the difference between the wave peak and the immediately following trough (local minimum). Wave latencies were defined as the time of each wave’s peak (maximum) relative to stimulus onset. In addition, as a readout for the magnitude of the entire ABR signal across all waves without making assumptions about changes in specific ABR waves, the RMS of the signal was calculated by applying the MATLAB function rms on the signal in the predefined response window, 1.6ms to 4ms, to include only the ABR signal. To account for longer response latencies at low sound levels, the response window was shifted by 0.16 ms (4 dps) for each 10 dB decrement.

Statistical comparisons of ABR wave-specific magnitudes and latencies were conducted for 90 dB SPL evoked ABRs. Assessed measures spanning ABRs across all sound levels were the RMS across waves I-V and wave II magnitude. Because of the potential disruption to the structure of the ABR after NOE and to ensure that equivalent waves were compared across conditions in each animal, the timing of wave II in the baseline conditions (BL) was used as time reference.

For the analysis of post NOE assessments, after applying the findpeaks function on the entire ABR signal, the closest peak to the reference time (wave II in BL) was defined as wave II peak. If two peaks were equally close to the latency of the BL wave II peak, the peak with the longer latency was selected. If wave II had been undefined in the BL condition, it was identified in the post NOE condition as described for the BL condition above (based on the maximum peak in a predefined detection window).

### Novel object sleep deprivation & EEG monitoring

Sleep deprivation for 6 hours (6hSD) was conducted via provision of novel objects in the home cage, which is an established method for inducing sleep deprivation in mice [e.g. 45,46], making use of the animals’ natural tendency to explore novel objects and therefore remain engaged. During the procedure, animals were monitored by the experimenter and novel objects were provided whenever the animal showed signs of drowsiness (behavioural immobility or increasing delta activity in the EEG, which was monitored while the animal was in the home cage). EEG/EMG data acquisition was performed using a Multichannel Neurophysiology Recording System (TDT). Cortical EEG was obtained from frontal and occipital derivations. EEG/EMG data were filtered between 0.1 and 100 Hz, amplified (PZ2 preamplifier, TDT) and visualised throughout the experiment.

Providing NOE during the last two hours of the animals’ active period (dark phase) within the 24h cycle (ZT 22-24) meant that the time immediately afterwards covered the period where sleep propensity was highest in all mice, ensuring a maximal difference in sleep amount between the two experimental groups. This timing was also chosen to take advantage of the fact that the effect of NOE is maximal during the active (dark) phase in mice: in Meltser et al. 2014 ^55^ NOE during the dark phase caused permanent hearing loss, whereas auditory thresholds were recovered by trKB-mediated protection against circadian sensitivity following noise overexposure during the light phase ^55^. Therefore, we optimised for both a maximal NOE effect in both groups as well as maximal sleep differences between groups.

### Statistical analysis

Statistical analysis of data from the GPIAS paradigm was performed using a generalised linear mixed model (GLMM, distribution: normal, link: identity) with the following multiple factors (repeated measures): experimental condition (sleep deprivation (6hSD) and control), time of testing (BL and Post NOE), order of presentation (time interval within assessments), response window (50 ms and 1000 ms durations) and, as random factor, animal identity, ID. Analysis was performed after exclusion of outliers. Outliers were defined as values larger than the mean of the bootstrapped raw data (5000 iterations) + 5 standard deviations.

For ABR recordings, statistical analysis of changes in magnitude and latency of the different waves (waves I-V) with stimuli at 90 dB SPL was performed using a GLMM on data with wave number (ABR wave I to V) as repeated factor and animals ID as random factor. Normality of data was assessed by inspection of QQ plots. Group differences in response modulation and response magnitude across stimulus levels were assessed using data after exclusion of outliers (values outside the mean ± 2 standard deviations) as input for a GLMM (repeated factors: experimental condition (6hSD and control), random factors: animal ID). Statistical analysis of ABRS was only performed on normalised data (Post NOE measure as % of BL).

EEG and EMG recordings were visually assessed to ensure efficiency of the sleep deprivation paradigm, but quantitative analysis was not conducted.

## Results

### Sleep deprivation after NOE mitigates gap detection impairment

After implantation of chronic electrodes and a recovery period (see Methods), baseline behaviour in the GPIAS paradigm was measured in all animals (Fig.2A). In line with previous studies ^43,56^, this paradigm served as the main tinnitus measure. GPIAS was assessed by comparing the startle response in gap-trials and non-gap trials, based on the assumption that in animals with normal hearing the startle response in gap trials is inhibited relative to non-gap trials (Fig.2B, see Methods for further details).

Under baseline conditions, before noise overexposure (NOE) and sleep deprivation, GPIAS ratios for both the control group (0.71±0.21, Fig.3A, BL) and the sleep deprivation (6hSD) group (0.8±0.21, Fig.3B, BL) were consistent with previously reported values in mice ^48,56–58^.

GPIAS values at baseline did not differ significantly between groups (*F*_*(1*,*210)*_*=3.24, p=0.07, n.s. Fig. 3C*). After noise exposure (NOE), the 6hSD group was kept awake for 6 hours, as confirmed through monitoring of each animal and EMG/EEG signals. Control animals, in contrast, slept during this period, as corroborated through subsequent sampling of video and EEG recordings.

**Figure 3.**
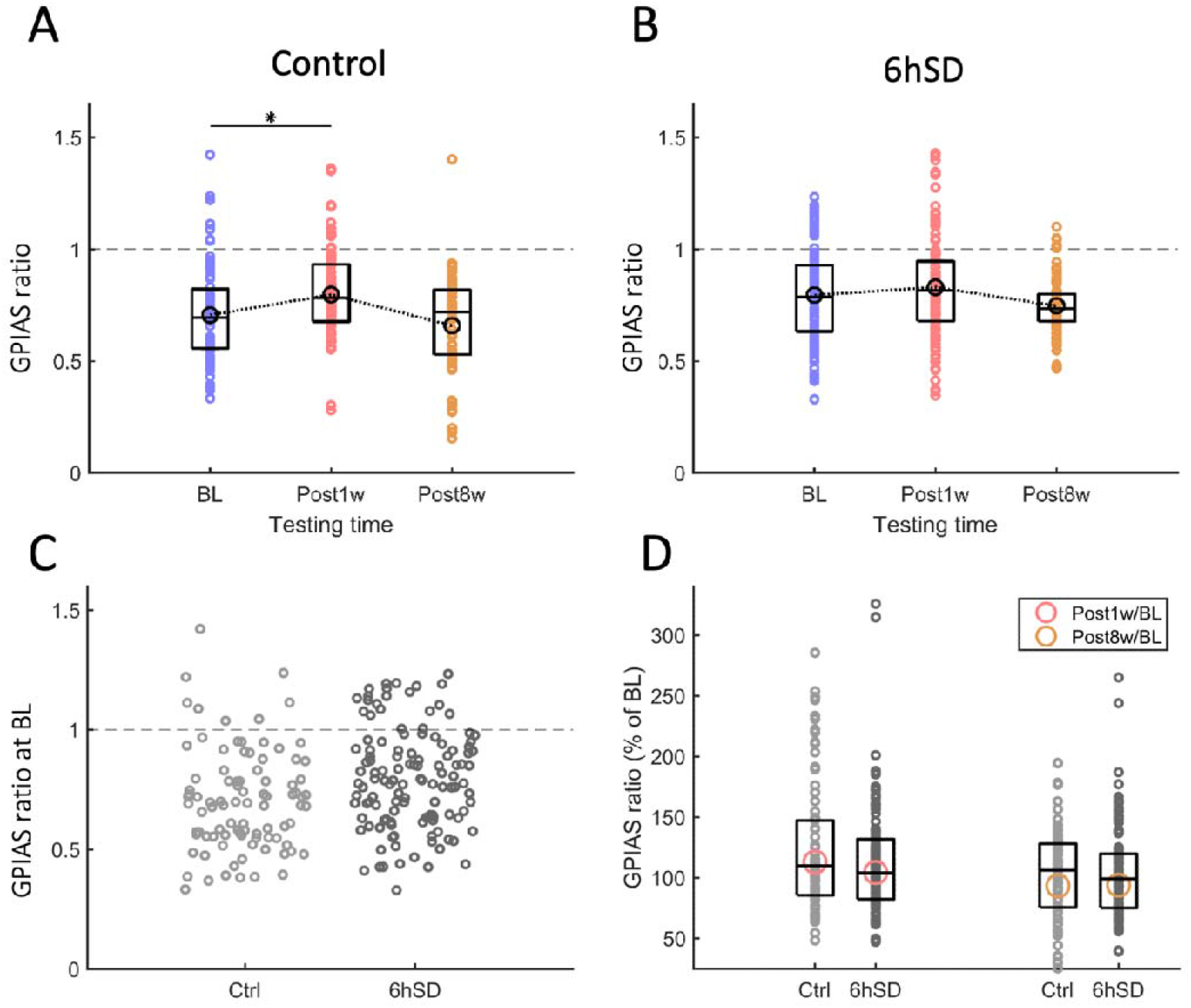
GPIAS ratio over time in 6hSD and control groups. **(A)** GPIAS ratio (startle magnitude in gap / startle magnitude in non-gap trials) separated by testing time. Datapoints depict individual values for the control group for both measures (peak and average response) and both windows (50 and 1000ms). Black circles overlaying the box blots depict mean values, connected by dotted lines. Box plots mark interquartile ranges, horizontal lines within box plots the median. The horizontal dashed line depicts GPIAS = 1, i.e. no gap induced startle inhibition. The horizontal solid line indicates a significant group difference with * indicating p<0.01, GLMM. **(B)** GPIAS ratio depicted as in A, but data are from the 6hSD group (6h novel object sleep deprivation after NOE). **(C)** GPIAS ratio at baseline for both Control (Ctrl, light grey) and sleep deprivation (6hSD, dark grey) groups. Datapoints for each group are dispersed across the x-axis for improved visibility. **(D)** GPIAS ratio as % of baseline. Data are shown for 1 week (red) and 6 weeks (right, red) post-NOE for control animals (Ctrl, light grey) and 6hSD animals (6hSD, dark grey). Coloured circles represent GPIAS as a percentage of baseline, calculated from the mean values for baseline (BL) and post-NOE conditions. Datapoints reflect changes in GPIAS between individual baseline and post-NOE values, where data are available for both timepoints. Missing datapoints (due to the absence of data for at least one of the timepoints) are 21 for the first plot, 42 for the second, 27 for the third, and 51 for the fourth.

**Figure 4.**
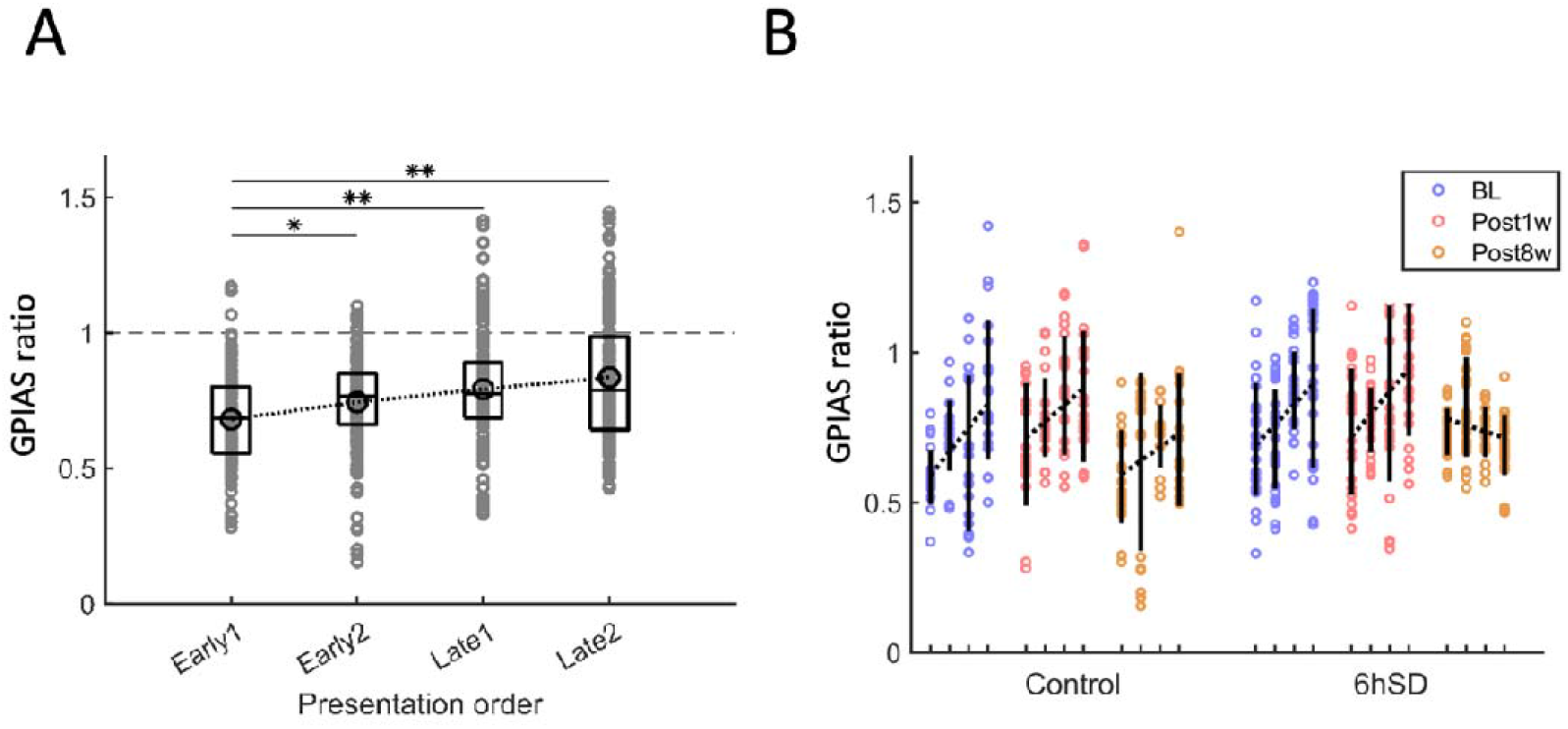
GPIAS: Intra-session dynamics in Control and 6hSD groups. **(A)** GPIAS ratio (= startle magnitude in gap trials / startle magnitude in non-gap trials) depicted across 4 consecutive intervals within the GPIAS assessment (presentation order): ‘Early1’, ‘Early2’, ‘Late1’ and ‘Late2’. For each interval, GPIAS was calculated based on 5 gap and 5 non-gap trials (‘Early1’: trials 1-5, ‘Early2’: trials 6-10, ‘Late1’: trials 11-15, ‘Late2’: trials 16-20 for gap and non-gap trials, respectively). Data points depict values of individual animals of both control and 6hSD groups, all testing times, measures (peak and average response) and windows (50ms and 1000ms). Black circles on top of the box plots depict mean values, connected by a dotted line across groups. Box plots mark interquartile ranges, horizontal lines within boxplots depict the median. The horizontal dashed line depicts GPIAS rate = 1, i.e. no gap induced startle inhibition. Horizontal solid lines indicate significant group differences with * indicating p<0.01 and ** p<0.001, GLMM. **(B)** GPIAS rate across 4 consecutive intervals (Early1, Early2, Late1, Late2) per assessment depicted as in panel A but separated by testing time (BL, Post1w, Post8w) and experimental group (Control and 6hSD). Error bars are standard deviations; black dotted lines are linear regressions across consecutive time intervals.

After NOE, both the control and the 6hSD group exhibited reduced gap detection ability, as indicated by higher GPIAS ratios compared to baseline (BL) (Fig. 3). As individual animals vary in their susceptibility to noise-induced tinnitus ^48,56–58^, we next examined within-group changes in addition to conducting a joint-group analysis.

There was no significant interaction between testing time (BL, 1 week post-NOE, 8 week post-NOE) and group (control and 6hSD) (F_(5,624)_ = 1.85, p = 0.10). Within-group analyses revealed a significant increase in GPIAS ratio in the control group (F_(2,271)_ = 4.20, p = 0.02; BL vs. Post1w: 0.71 ± 0.21 vs. 0.80 ± 0.21; β = 0.11, t_(273)_ = 2.64, p = 0.01; Fig. 3A), which was not evident in the 6hSD group (F_(2,353)_ = 1.47, p = 0.23; BL vs. Post1w: 0.80 ± 0.21 vs. 0.83 ± 0.23; β = 0.18, t_(355)_ = 0.36, p = 0.72; Fig. 3B). Indeed, the control group showed a 13% increase in GPIAS ratio relative to baseline (113% of BL), whereas the 6hSD group showed a smaller 4.4% increase (104.4% of BL; Fig. 3D, red markers).

Eight weeks after noise overexposure, GPIAS ratios were similar to BL levels in both groups (***Control***, *<u>BL vs Post8w</u>, 0.71±0.21 vs 0.66±0.21, β=0.02, t*_*(273)*_*=0.73*,*p=0.47*, ***<u>6hSD</u>***, *<u>BL vs Post8w</u>, 0.8±0.21 vs 0.75±0.12, β =-0.04, t*_*(355)*_*=-1.23, p=0.22*, Fig.3A,B), suggesting that the impairment was only temporary.

The transient impairment in GPIAS ratio in the control group was observed for both the startle peak and average magnitude (***Peak***, *F*_*(2*,*132)*_*=2.38, p=0.1, <u>BL vs Post1w</u>, β=0.1, t*_*(134)*_*=2.12, p=0.04, <u>BL vs Post8w</u>, β=0.02, t*_*(134)*_*=0.54, p=0.59*, ***Average***, *F*_*(2*,*136)*_*=5.75, p=0.004, <u>BL vs Post1w</u>, β=0.12*,*t*_*(138)*_*=2.69, p=0.01, <u>BL vs Post8w</u>, β=-6.677E-6, t*_*(138)*_*<0.001, p=1*). The transient impairment was also evident when only the long response window was used to define the startle magnitude (***1000ms window***, *F*_*(2*,*134)*_*=2.46, p=0.09, <u>BL vs Post1w</u>, β=0.1, t*_*(136)*_*=2.2, p=0.03, <u>BL vs Post8w</u>, β=-0.27, t*_*(136)*_*=-0.99, p=0.32*) and showed a trend towards significance for the short response window (***50ms window***, *F*_*(2*,*134)*_*=2.24, p=0.11, <u>BL vs Post1w</u>, β=0.11*,*t*_*(136)*_*=1.95*,*p=0.05, <u>BL vs Post8w</u>, β =-0.02, t*_*(136)*_*=-0.76, p=0.45*).

In the 6hSD group, GPIAS performance remained stable over time for all measures and response windows (***Peak***, *F*_*(2*,*175)*_*=2.11, p=0.12, <u>BL vs Post1w</u>, β=0.03, t*_*(177)*_*=0.66, p=0.51, <u>BL vs Post8w</u>, β=-0.04, t*_*(177)*_*=-1.21, p=0.23*, ***Average***, *F*_*(2*,*175)*_*=2.69, p=0.07, <u>BL vs Post1w</u>, β=-0.01, t*_*(177)*_*=-0.26, p=0.8, <u>BL vs Post8w</u>, β=-0.06, t*_*(177)*_*=-1.56, p=0.12*, ***50ms window, 1000ms window***, *F*_*(2*,*175)*_*=1.95, p=0.15, <u>BL vs Post1w</u>, β=-0.015*,*t*_*(177)*_*=-0.29, p=0.77, <u>BL vs Post8w</u>, β=-0.06, t*_*(177)*_*=-1.65, p=0.1*), except when GPIAS was assessed based on the short response window (50ms) where the 6HSD group showed a significant effect of time but no significant difference between BL and either Post1W or Post8W (Fig.3, *F*_*(2*,*175)*_*=3.57, p=0.03, <u>BL vs Post1w</u>, β=0.02, t*_*(177)*_*=0.42, p=0.67, <u>BL vs Post8w</u>, β=-0.04, t*_*(177)*_*=-1.62, p=0.11*,).

Therefore, the effect of sleep deprivation on transient GPIAS impairment after NOE is unlikely to be related to the way the startle response was measured. These findings should be interpreted cautiously given the absence of a statistically significant interaction, yet the comparison of individual group dynamics over time, as well as change scores are consistent with an attenuation of GPIAS impairment following sleep deprivation.

### Sleep deprivation following NOE affects GPIAS habituation

As previously shown ^59^ (revised in Wallace et al., 2024 ^60^), mice show habituation to the startle stimulus with a steady decrease in the startle amplitude within individual testing sessions and thus also less contrast between startle responses in gap and non-gap trials. In line with this, mice in this experiment showed a steady increase in the GPIAS ratio from the beginning to the end of testing sessions *(Effect of Interval, F*_*(3*,*626)*_*=10.49, p<0.001, Early1, 0.68±0.17, Early2, 0.75±0.18, Late1, 0.79±0.21, Late2, 0.84±0.24, Fig. 4A)*.

Next, we compared this intra-session habituation between early and late testing sessions (1 week versus 8 weeks after NOE) and found that this modulation of performance changed over time *(interaction between intra-session Interval (see Fig.4A, Testing time, F*_*(11*,*618)*_*=66.69, p<0.001)*. In the late assessment, 8 weeks after NOE, GPIAS values were more stable throughout the session than in earlier assessments and a smaller habituation effect was visible (Fig.4B). Notably, this was most apparent in the 6hSD group (see Fig.4B, 6hSD Post8W), suggesting that sleep deprivation following NOE may have long-term effects on the ability of animals to habituate to startling sounds.

In summary, these results suggest that NOE led to transiently impaired gap detection, a commonly used marker for tinnitus. Novel object sleep deprivation immediately after NOE mitigated this effect and even reduced the animals’ habituation to the startle stimulus within sessions, suggesting that both gap detection and sensitivity to startling stimuli may have been maintained or become heightened in these animals.

### Novel object sleep deprivation does not affect noise-induced auditory threshold changes

The effect of noise overexposure on central auditory function was estimated using ABRs (Fig. 5A). ABR threshold assessment as well as automated analysis of ABR waves I to V (Fig. 5B & see Methods) were conducted at baseline conditions and 1 week after NOE.

**Figure 5.**
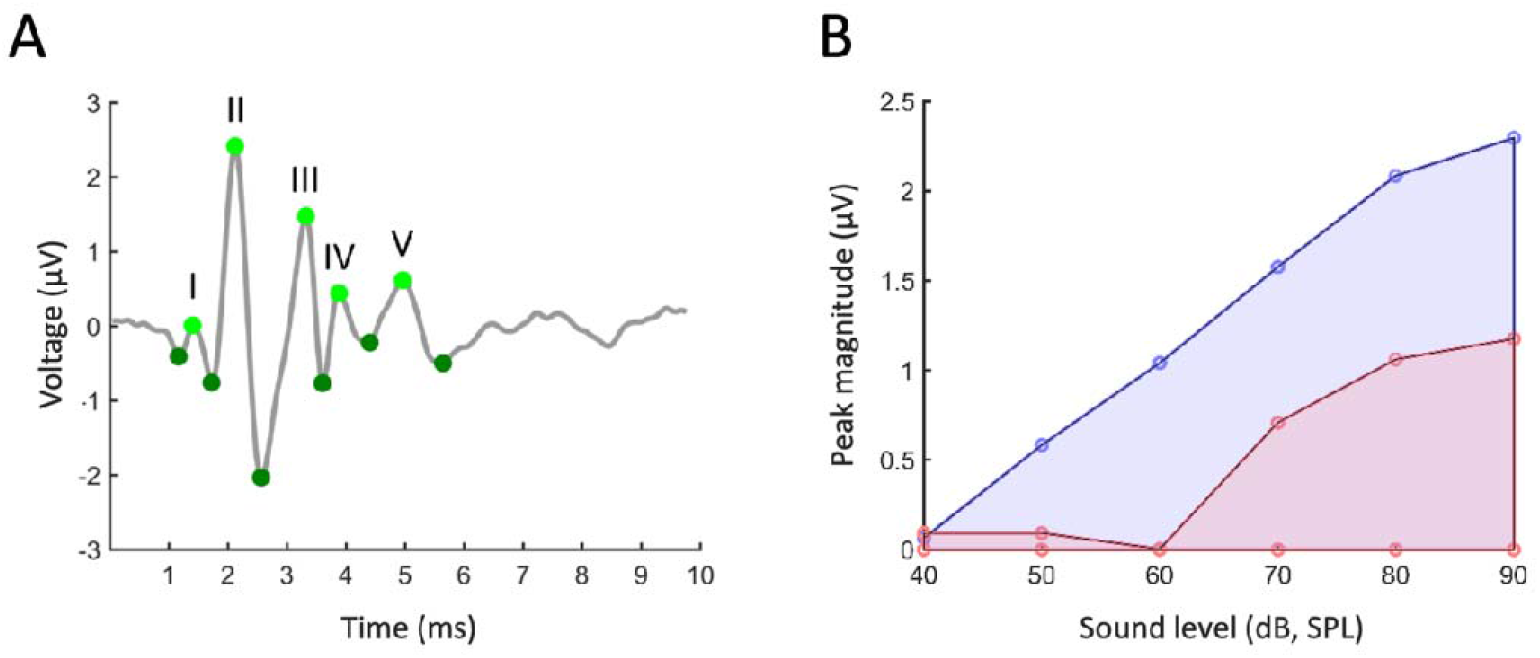
Auditory brainstem responses (ABRs): Methodology and analysis. **(A)** Example of an ABR signal to 90dB SPL click stimuli (average over 700 trials). Green markers depict peaks and throughs of waves (I to V) as detected with a custom-written MATLAB algorithm. **(B)** The magnitude of the ABR response across all sound intensities was estimated by calculating the area under the level-response graph (here the area enclosed by the blue ‘BL’ and red ‘post NOE’ graph, respectively).

Pure tone ABR thresholds decreased with increasing sound frequency over the range of values tested (*effect of sound frequency*, <u>*F*_*(3*,*162)*_*=258.85, p<0.001)*</u>. The lowest thresholds were measured for clicks (Fig. 6). To assess threshold changes over time, both groups were compared to a joint baseline. There was an overall effect of time on measured thresholds for both the control group (*F*_*(1*,*118)*_*=66.1, p<0.001*) and the 6hSD group (*F*_*(1*,*130)*_ *= 186.63, p<0.001*), suggesting that noise overexposure led to a threshold change. However, this change was not significant for any individual stimulus: *w*hen comparing thresholds before and after NOE, for both the control group and 6hSD group, thresholds for 1 kHz pure tones were constant across timepoints (Fig.6, 1 kHz). Raw p-values from the remaining stimuli were adjusted for multiple comparisons using the Holm (sequential Bonferroni) method. After correction, none of the frequencies showed significant differences between baseline and post NOE assessment for either the control group (Holm adjusted p-values: 0.15, 0.84, 0.84, 0.84, 0.84 for 2, 4, 8,16 kHz pure tones and clicks, respectively) or for the 6hSD group (Holm adjusted p-values: 0.75, 0.75, 0.87, 0.87, 0.87), indicating that noise-induced auditory threshold changes were not affected by acute sleep deprivation after NOE.

**Figure 6.**
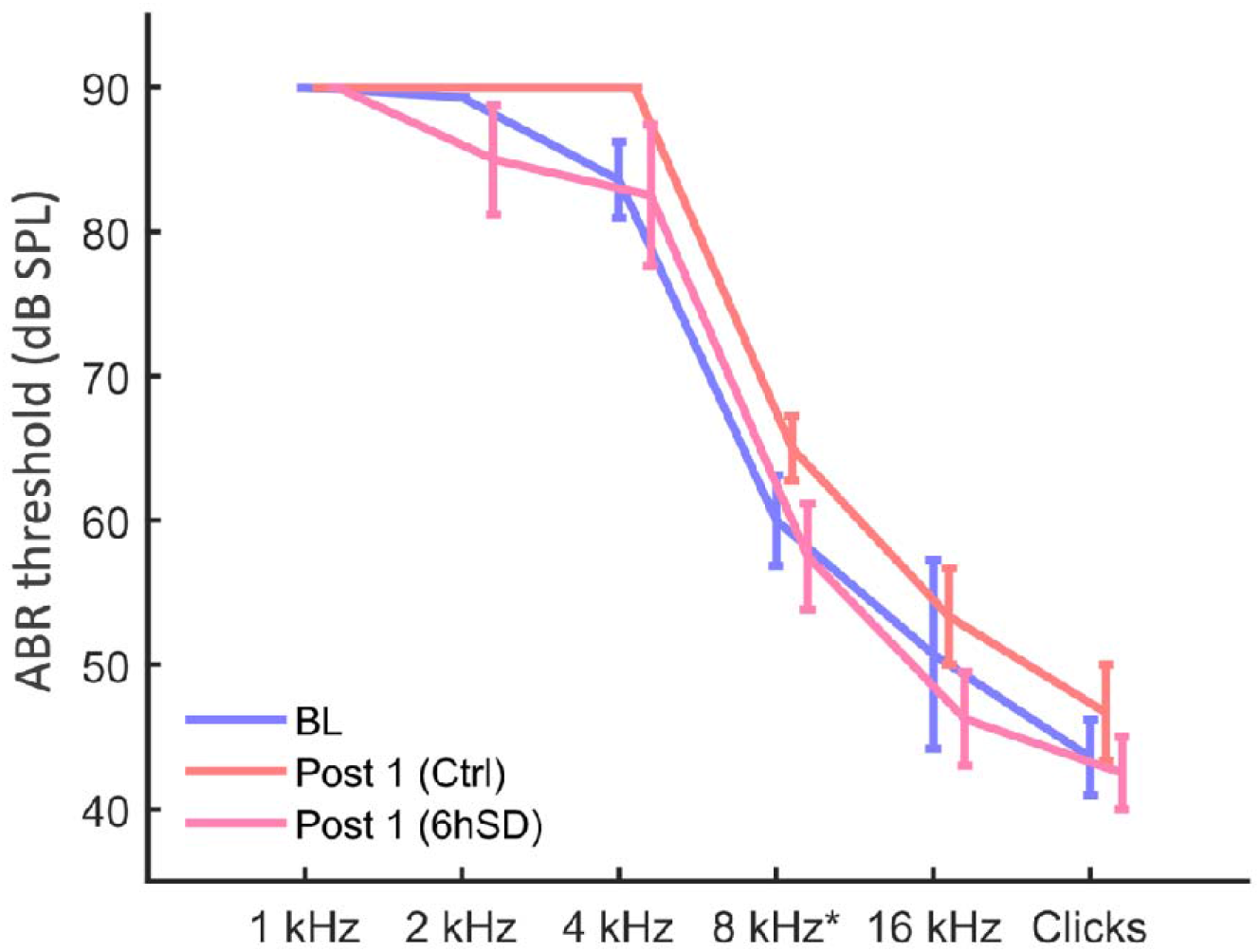
Auditory brainstem response (ABR) thresholds. Thresholds for control and 6hSD groups before and after NOE. Thresholds were determined through visual inspection of raw data under randomised and blind conditions. Data are presented as means ± standard errors for all measured stimuli. Discrete threshold values, ranging from 30 to 90 dB SPL in 10 dB steps, were assigned. Due to the limited range of distinct values, the error bars are absent (standard error = 0) for 1, 2 and 4 kHz stimuli in some cases.

### Sleep deprivation following NOE does not affect click-evoked ABR magnitude and latency

To assess in more detail how noise overexposure impacted auditory sensitivity, ABR response magnitudes and latencies were assessed, initially focusing on click stimuli. Animals showed distinct wave magnitudes across ABR waves I to V in response to 90 dB SPL click stimuli (Fig 5A and 7A), which facilitated automated wave detection (see Methods: Automated ABR signal detection). To assess the direction and magnitude of changes in individual ABR waves following NOE, the change in each respective measure was defined as:

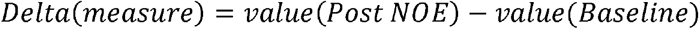

**Figure 7.**
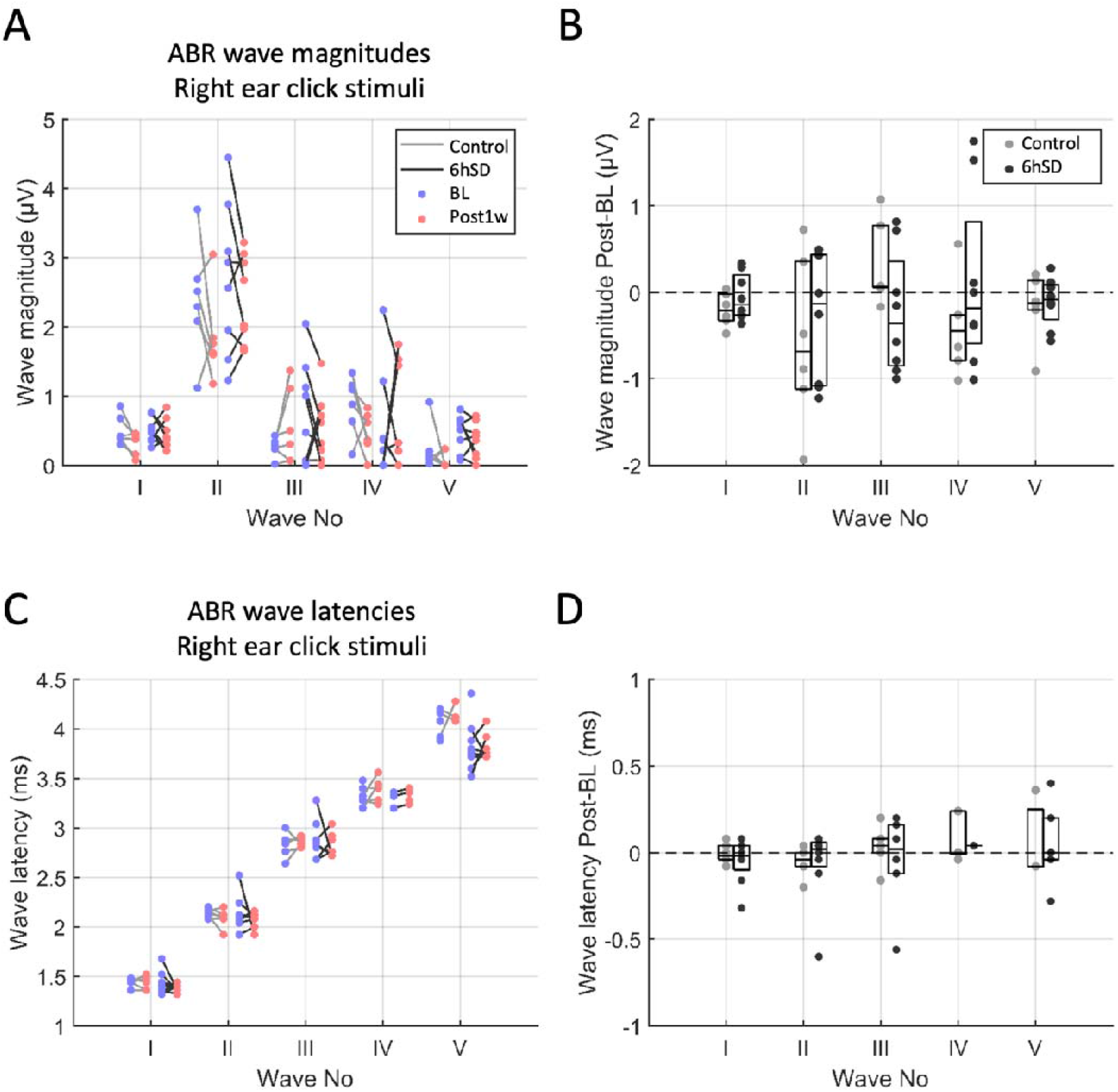
Auditory brainstem responses (ABRs) for 90 dB SPL click stimuli: individual waves. **(A)** Magnitude of individual ABR waves (I-V) before and after NOE for both Control and 6hSD groups (see figure key). Datapoints are animal averages. Lines connect animal averages obtained during the BL and Post1w assessments. **(B)** Difference in wave magnitude (Post – BL values) calculated for each animal. Data points represent individual animals. Box plots mark interquartile ranges, horizontal lines within boxplots depict the median. Values below 0 (dashed line) depict a decrease in wave magnitude after NOE relative to BL. **(C)** Wave latencies, shown as in A. **(D)** Wave latency changes from BL to Post1w, shown as in **B**.

Animals tended to show reduced response magnitudes especially in ABR wave I *(<u>Controls</u>, −0.2±0.19µV, <u>6hSD</u>, −0.06±0*.27µV) and wave II *(<u>Controls</u>, −0.56±0.98µV, <u>6hSD</u>, −0.29±0.74µV)* following NOE (Fig.7B). Interestingly, control animals showed an elevation of wave III magnitude (*by 0.31±0.49µV relative to BL*), which was not apparent in 6hSD animals *(−0.24±0.71µV*), though there was no significant difference in wave magnitude changes following NOE between control and 6hSD animals for any ABR wave (*interaction ABR wave identity and group, F*_*(9*,*60)*_*=1.33, p=0.24*, Fig.7B). Regarding response latencies (Fig.7C,) there was an interaction between wave identity and group (*F*_*(9*,*48)*_*=3.99, p<0.001*), but latencies did not show significant difference between groups for any specific wave identity (***Wave I latency***, *<u>Control vs 6hSD</u>, −0.01±0.06 vs −0.05±0.13ms, F*_*(1*,*12)*_*=0.76, p=0.4*, ***Wave II latency***, *<u>Control vs 6hSD</u>, −0.05±0.09 vs −0.07±0.23ms, F*_*(1*,*12)*_*=2.36, p=0.15*, ***Wave III latency***, *0.03±0.12 vs −0.05±0.28ms, F*_*(1*,*10)*_*=0.08, p=0.79, <u>Control vs SD</u>*, ***Wave IV latency***, *<u>Control vs 6hSD</u>, 0.09±0.14 vs 0.04±0.0ms, F*_*(1*,*6)*_*=2.36, p=0.18*, ***Wave V latency***, *<u>Control vs 6hSD</u>, 0.07±0.25 vs 0.06±0.22, F*_*(1*,*8)*_*=1.47, p=0.26*, Fig.7D). This suggests that sleep deprivation did not change the effect of noise overexposure on high-intensity click-evoked responses, in line with thresholds observed for click train stimuli (Fig.6).

To assess whether sleep deprivation influenced the effect of NOE on the total ABR magnitude, calculated across responses to all tested stimulus levels (40, 50, 60, 70, 80, 90 dB SPL clicks), the area under the level-response curve was calculated for each animal and assessment (Fig. 5B). The level-response curve was produced for two response measures, the amplitude of ABR Wave II and the RMS amplitude of the signal, integrating the overall magnitude of ABR Waves I–V.

Irrespective of the experimental group, animals showed lower total ABR magnitudes following NOE as compared to BL for ABR wave II (<u>control</u>: *74.01±41.64 % of BL and <u>6hSD</u>: 94.88±27.89% of*) and the RMS (<u>control</u>: *80.22±31.86% of BL and <u>6hSD</u>: 81.53±10.99% of BL*), suggesting that total ABR response levels were affected by NOE in both groups (Fig.8A). The decrease in the total magnitude of wave II tended to be more pronounced in control animals than in the 6hSD group (***Wave II total magnitude***, *<u>Control vs 6hSD</u>, 74.01±41.64 vs 94.88±27.89% of BL, F*_*(1*,*12)*_*=4.58, p=0.05*, Fig.8A), indicating that, measured across stimulus levels, the ABR response was more impaired following noise overexposure when animals were allowed to sleep in the subsequent six hours. The decrease in total response magnitude across all ABR waves was similar between control and 6hSD animals (***RMS total magnitude***, *<u>Control vs 6hSD</u>, 80.22±31.86 vs 81.53±10.99% of BL, F*_*(1*,*11)*_*=0.02, p=0.9*, Fig.8A).

**Figure 8.**
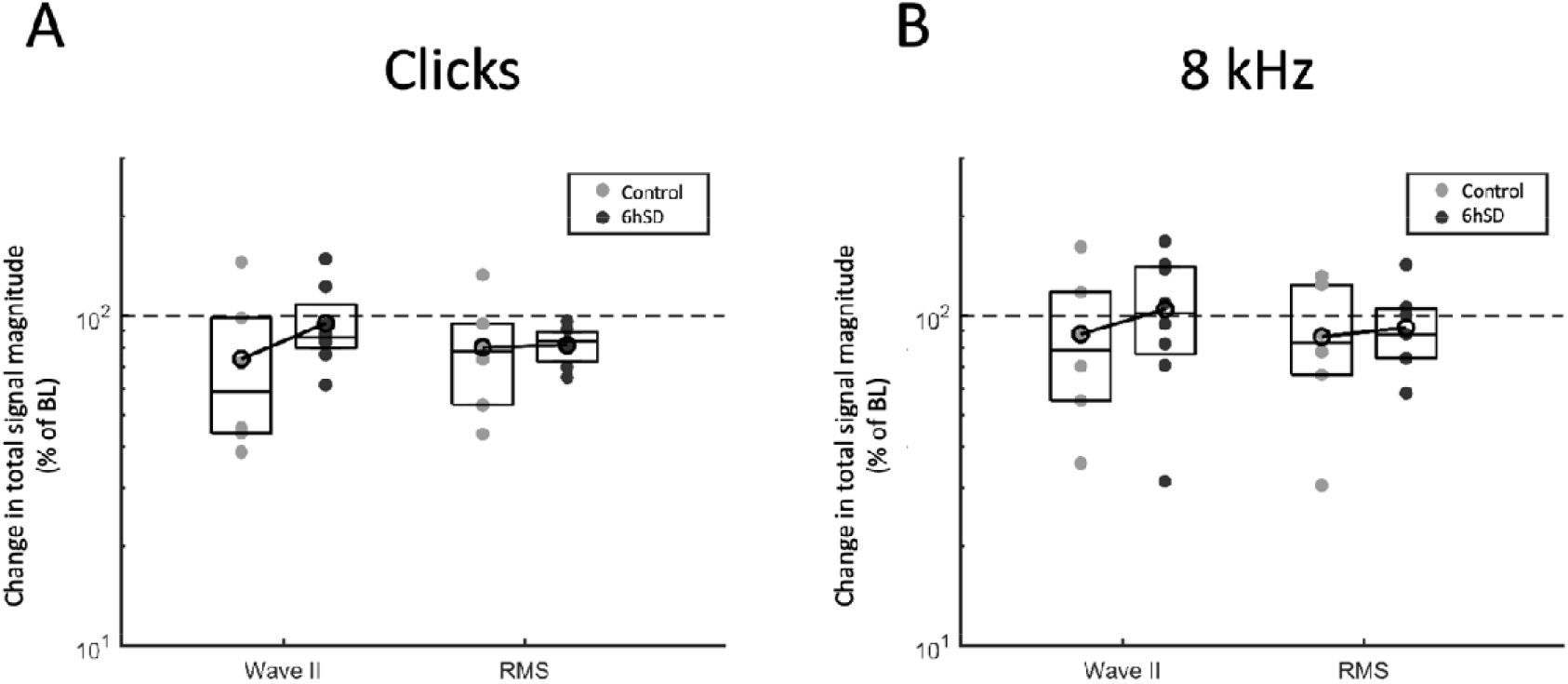
Auditory brainstem responses (ABRs) for click and 8 kHz stimuli. Changes after NOE in total magnitude in Control and 6hSD groups. ABR total magnitude (area under the level-response graph, as % of BL), shown for ABR wave II and the RMS of the signal for click stimuli **(A)** and 8 kHz stimuli **(B)**. Box plots mark interquartile ranges, horizontal lines within boxplots depict the median. Black circles on top of the box plots depict mean values, connected by lines across groups. Values above 1 represent an increase in the respective measure, values below 1 (dashed line) represent a decrease relative to the BL assessment.

Thus, while click-evoked ABRs tended to decrease after NOE in all animals and sleep deprivation was associated with less impairment, no significant difference between sleep deprived and control groups were evident.

### Sleep deprivation after NOE reduces gain elevation for 8kHz pure tones

Since noise overexposure was conducted using an 8 kHz centred NBN stimulus, hearing impairments were expected to be most prominent for stimuli in a similar frequency range to 8Khz, as also suggested by previous studies in animal models of noise-induced hearing loss ^61–63^. Therefore, ABRs for 8 kHz pure tones were assessed in more detail, based on response magnitudes and latencies.

Responses to 90 dB SPL tone stimuli had overall lower magnitudes than those evoked by click stimuli (compare Fig.7A & Fig.9A, note different y-axes dimensions). However, wave II was still the most pronounced component of the ABR signal and waves were clearly discernible based on their magnitude (Fig.9A).

**Figure 9.**
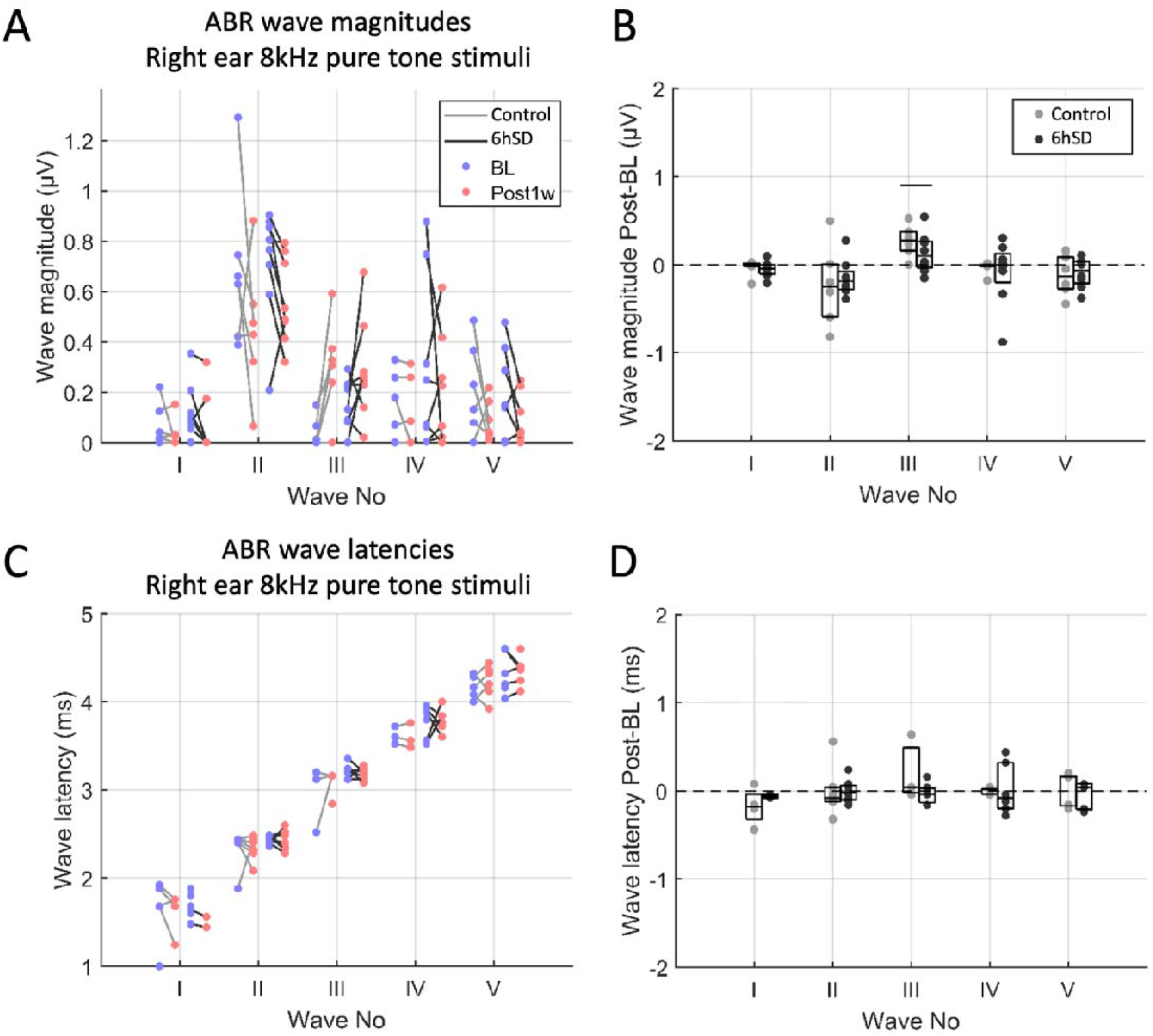
Auditory brainstem responses (ABRs) for 90 dB SPL 8kHz pure tone stimuli: individual waves. (A) Magnitude of individual ABR waves (I-V) before and after NOE for both Control and SD groups (see figure key). Datapoints are animal averages. Lines connect animal averages obtained during the BL and Post1w assessments. (B) Difference in wave magnitude (Post – BL values) calculated for each animal. Data points represent individual animals. Box plots mark interquartile ranges, horizontal lines within boxplots depict the median. Values below 0 (dashed line) depict a decrease in wave magnitude after NOE relative to BL. The horizontal line depicts a statistically significant group difference (p<0.05). (C) Wave latencies, shown as in A. (D) Wave latency changes from BL to Post1w, shown as in B.

Following noise overexposure, the magnitudes of wave I and II tended to be lower than in the baseline assessment in both experimental groups (***Wave I magnitude***, *<u>Controls</u>, −0.04±0.9µV, <u>SD</u>, −0.05+-0.9µV*, ***Wave II magnitude***, *<u>Controls</u>, −0.24±0.46µV, <u>6hSD</u>, −0.15±0.21µV*, Fig.9B), whereas wave III was enhanced (***Wave III magnitude***, *<u>Controls</u>, 0.27±0.18µV, *6hSD*, 0.13±0.23µV*, Fig.9B). Unlike for click stimulation, differences between control and 6hSD animals were evident in ABR wave-specific changes in magnitude, as indicated by a strong interaction between ABR wave identity and experimental group (*F*_*(9*,*16)*_*=9.7, p<0.001*): in particular the elevation in magnitude of ABR wave III was significantly more pronounced in control than in 6hSD animals (*F*_*(1*,*12)*_*=7.85, p=0.02*, Fig.9B). This indicates that NOE may have led to reduced evoked activity in early auditory processing stages (auditory nerve and cochlear nucleus) and to a compensatory gain increase in upstream regions associated with ABR wave III, such as in the superior olivary nucleus ^64^, which was especially pronounced in control animals. Note that a similar trend was visible for click stimulation in control animals (Fig.7B). In other tone-evoked ABR waves, the changes after NOE were comparable between control and SD groups (***Wave I magnitude***, *<u>Control vs 6hSD</u>, −0.04±0.09 vs −0.05±0.09µV, F*_*(1*,*12)*_*=0.19, p=0.67*, ***Wave II magnitude***, *<u>Control vs 6hSD</u>, −0.24±0.46 vs −0.15±0.21µV, F*_*(1*,*12)*_*=0.52, P=0.49*, ***Wave IV magnitude***, *<u>Control vs 6hSD</u>, −0.03±0.7 vs −0.09±0.37µV, F*_*(1*,*12)*_*=0.08, p=0.78*, ***Wave V magnitude***, *<u>Control vs 6hSD</u>, −0.12±0.23 vs −0.09±0.17µV, F*_*(1*,*12)*_*=1.56, p=0.24*).

Although there was an interaction between latency changes for 90 dB SPL stimuli and group (*F*_*(9*,*39)*_*=38.85, p<0.001*), this was not evident for any particular ABR wave (Fig.9C,D).

The total ABR signal magnitude (encompassing responses to all stimulus levels) was similarly affected in control and 6hSD animals, even though total magnitude of wave II tended to be more impaired in control animals (***Wave II total magnitude***, *<u>Control vs 6hSD</u>, 87.99±45.62 vs 104.72±44.34% of BL, F*_*(1*,*12)*_*= 3.77, p=0.08*, Fig.8B). As with click-evoked ABRs, this was not evident for total magnitude of the RMS encompassing ABR waves I to V (***RMS total magnitude***, *<u>Control vs 6hSD</u>, 86.48±37.7 vs 92.13±27.82% of BL, F*_*(1*,*11)*_*=0.29, P=0.6*, Fig.8B).

Overall, 8 kHz pure tone evoked ABRs tended to show larger post-NOE impairment in control than in 6hSD animals, although this was not a significant effect, consistent with the results based on click stimuli. Interestingly, control animals showed pronounced elevation of ABR wave III, reminiscent of hyperexcitability, and significantly more so than 6hSD animals. This further supports the notion that sleep deprivation may be a mitigating factor for the consequences of noise overexposure.

## Discussion

The aim of this study was to assess whether sleep plays a role in generating tinnitus following overexposure to noise. To address this, a group of mice (n=8) was kept awake for six hours (6hSD) immediately after noise exposure. Evidence for tinnitus and hearing loss was assessed and compared to a group of control mice (n=6) that was left undisturbed following NOE.

### Sleep deprivation mitigates effects of noise trauma

After NOE, all animals showed signs of experiencing tinnitus, indicated by the transiently lower gap detection ability as assessed by the change in GPIAS scores, but this impairment was significant only in the control group. This suggests that the novel object sleep deprivation protocol after NOE may have reduced the noise-induced impairment of gap detection, likely due to less tinnitus or hearing loss compared to animals that were allowed to sleep.

Further to maintaining their gap detection scores, 6hSD animals showed, in the long term, less habituation within sessions than control animals. This suggests that 6hSD animals remained sensitive to the startle stimulus throughout the assessment. This indicates that they not only maintained their gap detection responses after NOE, but that they may eventually have developed increased sensitivity to startle sounds, thereby decreasing the rate of habituation after repeated exposure within assessments. This could be connected to a latent development of hyperacusis, which is thought to develop over time due to deafferentation following noise overexposure ^49^.

To assess whether NOE caused a hearing impairment, ABRs were acquired for all animals before and after NOE. Reduced responses, particularly for wave II and the ABR RMS, were apparent for all animals after NOE, for both click stimuli and 8 kHz pure tones. It is unlikely, however, that this observed hearing impairment could explain the group-specific deficits in the GPIAS following NOE, as animals from both groups still showed marked and comparable startle responses on no-gap trials (Supp.Fig.1). Control animals also showed signs of elevated activity reflected in ABR wave III (corresponding to activity in the superior olivary nucleus ^64^) following NOE, especially for pure tone stimulation using the centre frequency of the NOE stimulus (8 kHz). This may indicate increased synchrony or hyperexcitability, which is often described as a functional correlate of noise-induced tinnitus ^8,9,14,46,65,66^. Such activity changes have been described at various stages of the ascending auditory pathway, including the cochlear nucleus ^5,6,46^ and the inferior colliculus ^9,10,67^. Moreover, the superior olivary nucleus has specifically been shown to mediate hyperresponsiveness in tinnitus sufferers ^68^, suggesting that the elevated ABR wave III magnitude observed in control animals in this study might be connected to tinnitus.

In summary, a reduction in GPIAS values and the wave-specific changes in ABR magnitude were all more pronounced in control than in 6hSD animals while ABR thresholds did not change in either group after NOE. This suggests that novel object sleep deprivation did not affect noise-induced hearing loss but mitigated tinnitus, both behaviourally and physiologically, and may therefore have a protective effect against the consequences of noise exposure.

### Design and characteristics of the sleep deprivation paradigm

To ascertain what aspects of the experimental design may underly the observed group differences, it is important to clarify the rationale behind the timing and design of the used sleep deprivation paradigm.

Acute sleep deprivation rather than chronic sleep restriction was chosen for this experiment as the former is associated with only brief impairments in attention and cognitive abilities, which return to baseline levels after only one night of recovery sleep in human subjects ^69^. Acute sleep deprivation also limits the amount of potentially stressful experience that may be predisposing for tinnitus ^70^ and therefore minimises confounding effects of the procedure. Furthermore, sleep-dependent plasticity triggered by waking experience is known to already take place in the first night of sleep after learning a task ^30^. Ocular dominance plasticity after monocular occlusion has also been shown to occur in the first six hours of the subsequent sleep period ^28,29^, suggesting that sleep-dependent plasticity can occur or start within a short timeframe after waking experience and is sensitive to acute sleep deprivation within that time. Sleep-dependent plasticity involved in tinnitus development might therefore also be affected by acute sleep deprivation soon after noise trauma.

We followed an established protocol in which novel objects were used to keep animals engaged and awake(e.g. ^44,54^). This method of sleep deprivation is based on the animal’s tendency to explore changes in its environment when novel objects are introduced into the cage, and is thought to minimise procedure-related stress relative to more intrusive approaches, such as handling. Importantly, any stress induced by the sleep deprivation paradigm is unlikely to explain the effects observed in the 6hSD animals. Since stress is a known predisposing factor for tinnitus development ^70^, any sleep deprivation-related stress should have led to results opposite to what we observed. That said, increased exploratory behaviour linked to novel object sleep deprivation may alter the auditory environment compared to control mice. This is relevant, as acoustic stimulation after noise overexposure can help mitigate tinnitus or hearing loss [61,66]. However, care was taken to minimise environmental noise during the sleep deprivation paradigm: both groups were kept in the same room, and sleep deprivation only involved placing and removing small objects by a single experimenter, without using acoustic stimulation to keep animals awake. While the procedure and the animals’ exploratory behaviour might have provided some acoustic enrichment, it differs from previous studies where prolonged acoustic enrichment with stimuli specifically matched to the hearing loss range helped mitigate noise trauma [61,66]. Therefore, the difference in sleep opportunity after NOE is likely to be the most relevant factor distinguishing control and 6hSD animals.

### Multiple attributes of sleep may mediate consequences of noise trauma

The fact that six hours of acute sleep deprivation after NOE were associated with reduced tinnitus symptoms suggests that sleep may mediate or amplify such impairments. Several sleep-related factors could contribute to this effect.

Sensory disconnection during sleep ^71^ plays a role in shielding the animal from an acoustic environment. It has previously been shown that acoustic enrichment after noise trauma mitigates symptoms of hearing loss and tinnitus ^66,72^. It is of note, however, that auditory evoked activity is largely maintained during sleep and there are indications that sensory disconnection takes place only at the cortical level ^73^. Thus, sleep does not fully prevent the acoustic environment from shaping brain activity ^74^, particularly at the brainstem level, even though this may be different from the waking state ^75^.

Moreover, sleep is involved in mediating and consolidating a range of plastic processes that may be relevant to tinnitus development ^28–33^, such as cortical map plasticity ^29,76^ and other systems level plasticity ^30^. It is possible that sleep contributes to tonotopic map plasticity, which has been implicated in tinnitus development ^12,77^ and is associated with hearing loss ^78^. Changes in non-auditory brain regions, possibly mediated by sleep-dependent systems-level plasticity, are thought to underlie the establishment of a brain-wide and limbic representation of tinnitus ^19,21^. Yet sleep-dependent plasticity underlying persistent tinnitus likely extends beyond these examples. For instance, the scaling of synapses ^36^ or spatially restricted synapse plasticity ^34^ during sleep might add to the number of processes involved in tinnitus formation.

Although our study addressed the effect of sleep immediately after NOE, it is important to acknowledge a possible effect of sleep-wake history on auditory function. Meltser et al. (2014)^55^ reported increased susceptibility to NOE during the dark phase in mice, suggesting potential roles for circadian timing and prior sleep history. More recent studies further showed that sleep manipulation alters auditory function and resilience to noise trauma. For example, three weeks of sleep fragmentation in mice reduced ABR wave I amplitude and increased ABR latency^79^, indicating that sleep fragmentation alone can impair auditory sensitivity. Another study showed that five days of REM sleep deprivation before NOE can exacerbate cochlear damage, based on ABR thresholds and cochlear synaptic ribbon counts^80^. Interestingly, the same study showed that only 24 h of REM sleep deprivation reduced cochlear damage and suggested that an adaptive autophagy response to acute stress may underlie this protective effect^80^. Our sleep manipulation differed substantially from these approaches, as we applied exploration-based novel object sleep deprivation, which keeps animals awake and engaged, instead of fragmenting their sleep, and applied it after rather than before NOE. This design allowed us to specifically address the role of sleep in tinnitus development following a triggering event. Nevertheless, it remains possible that an adaptive stress response also contributed to the SD-induced resilience observed in our study.

The negative effect of tinnitus on sleep has long been acknowledged, such as its potential to disturb and disrupt sleep ^81–88^. Less focus has been placed on investigating the effect of sleep, or lack thereof, on tinnitus ^18,89–91^. Similarly, the likely bidirectional relationship between tinnitus und sleep ^41^ and how sleep may be harnessed to treat tinnitus ^90,90^ has received little attention thus far. Recently, we described in a ferret model of chronic tinnitus the association between tinnitus and sleep disruption ^50^ and demonstrated that neural markers of tinnitus were reduced during sleep, suggesting that sleep may transiently alleviate tinnitus. Our current results obtained in the mouse tinnitus model indicate that sleep immediately following NOE may provide a critical time window that could be targeted to reduce the consequences of noise exposure even before they arise.

While future studies will be needed to determine which aspect of sleep can affect the development and consolidation of noise-induced hearing impairments and tinnitus, the results of this study suggest that modulation of natural brain state dynamics provide fruitful ground for mitigating the consequences of noise overexposure. More broadly, our results add an important facet to the growing body of evidence that sleep and wake dynamics not only reflect neural or sensory pathology but may also represent a target for clinical intervention in hearing-related disorders.

## Supplementary data

**Supp. Fig. 1:**
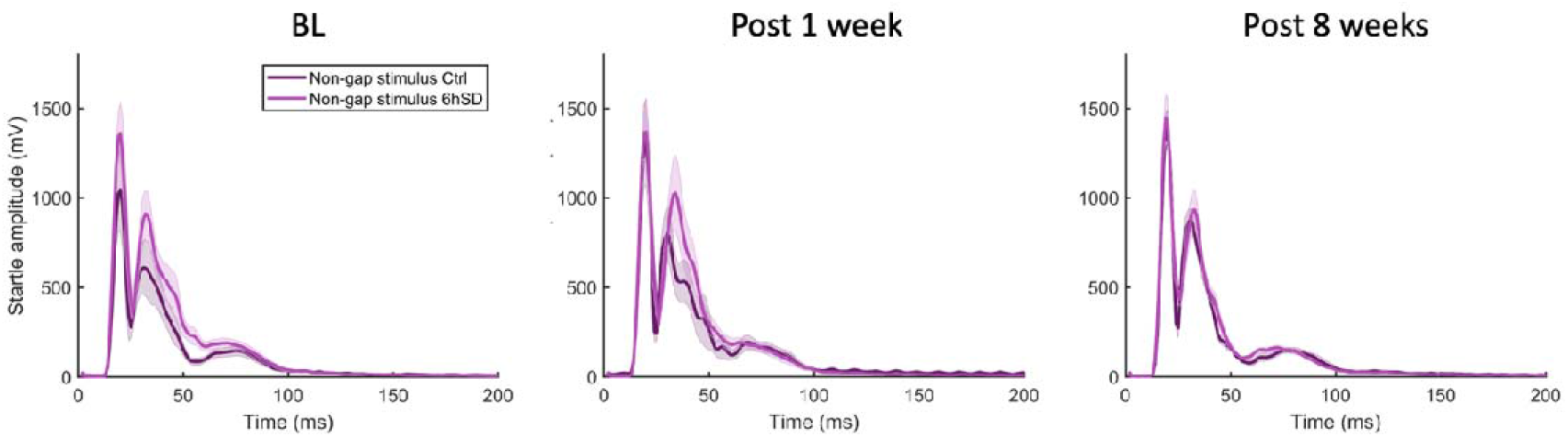
Startle responses to non-gap stimuli across time. Average startle signal in non-gap trials, i.e. trials where the startle sound was not pre-empted by a silent gap in the background noise, for animals in the control group (dark purple) and in the 6hSD group (bright purple). Startle signals are shown for baseline (BL, left panel), 1 week after NOE (Post 1 week, middle panel) and 8 weeks after Noe, (Post 8 weeks, right panel). Shading represents standard errors.

**Suppl. Table 1:** Auditory brainstem response (ABR) thresholds. Thresholds for individual animals in the control group (n=6) and 6hSD group (n=8) before and after NOE. **Top:** joint BL for Ctrl and SD groups. **Middle**: post NOE assessment for the control group. **Bottom**: post NOE assessment for the 6hSD group. Thresholds were determined through visual inspection of raw data under randomised and blind conditions.

|  | AnimalNumber | 1 kHz | 2 kHz | 4 kHz | 8 kHz* | 16 kHz | Clicks |
| --- | --- | --- | --- | --- | --- | --- | --- |
| 1 | 1 | 90 | 90 | 90 | 50 | 70 | 40 |
| 2 | 2 | 90 | 90 | 90 | 70 | 60 | 50 |
| 3 | 3 | 90 | 80 | 70 | 50 | 50 | 40 |
| 4 | 4 | 90 | 90 | 70 | 60 | 50 | 50 |
| 5 | 5 | 90 | 90 | 80 | 60 | 50 | 40 |
| 6 | 6 | 90 | 90 | 80 | 50 | 60 | 40 |
| 7 | 7 | 90 | 90 | 80 | 60 | 30 | 40 |
| 8 | 8 | 90 | 90 | 90 | 70 | 80 | 40 |
| 9 | 9 | 90 | 90 | 90 | 50 | 40 | 40 |
| 10 | 10 | 90 | 90 | 90 | 70 | 30 | 40 |
| 11 | 11 | 90 | 90 | 90 | 70 | 30 | 40 |
| 12 | 12 | 90 | 90 | 70 | 70 | 60 | 60 |
| 13 | 13 | 90 | 90 | 90 | 60 | 40 | 40 |
| 14 | 14 | 90 | 90 | 90 | 50 | 60 | 50 |
| 1 | 1 | 90 | 90 | 90 | 60 | 50 | 50 |
| 2 | 2 | 90 | 90 | 90 | 60 | 60 | 40 |
| 3 | 3 | 90 | 90 | 90 | 70 | 60 | 60 |
| 4 | 4 | 90 | 90 | 90 | 70 | 50 | 40 |
| 5 | 5 | 90 | 90 | 90 | 60 | 40 | 40 |
| 6 | 6 | 90 | 90 | 90 | 70 | 60 | 50 |
| 1 | 1 | 90 | 90 | 90 | 50 | 60 | 40 |
| 2 | 2 | 90 | 90 | 50 | 60 | 60 | 40 |
| 3 | 3 | 90 | 90 | 90 | 60 | 40 | 40 |
| 4 | 4 | 90 | 90 | 90 | 50 | 40 | 40 |
| 5 | 5 | 90 | 90 | 90 | 60 | 40 | 60 |
| 6 | 6 | 90 | 60 | 90 | 50 | 40 | 40 |
| 7 | 7 | 90 | 90 | 80 | 80 | 40 | 40 |
| 8 | 8 | 90 | 80 | 80 | 50 | 50 | 40 |

